# A variant by any name: quantifying annotation discordance across tools and clinical databases

**DOI:** 10.1101/054023

**Authors:** Jennifer Yen, Sarah Garcia, Aldrin Montana, Jason Harris, Steven Chervitz, John West, Richard Chen, Deanna M. Church

## Abstract

**Background:** Clinical genomic testing is dependent on the robust identification and reporting of variant-level information in relation to disease. With the shift to high-throughput sequencing, a major challenge for clinical diagnostics is the cross-identification of variants called on their genomic position to resources that rely on transcript- or protein-based descriptions.

**Methods:** We evaluated the accuracy of three tools (SnpEff, Variant Effect Predictor and Variation Reporter) that generate transcript and protein-based variant nomenclature from genomic coordinates according to guidelines by the Human Genome Variation Society (HGVS). Our evaluation was based on comparisons to a manually curated list of 127 test variants of various types drawn from data sources, each with HGVS-compliant transcript and protein descriptors. We further evaluated the concordance between annotations generated by Snpeff and Variant Effect Predictor with those in major germline and cancer databases: ClinVar and COSMIC, respectively.

**Results:** We find that there is substantial discordance between the annotation tools and databases in the description of insertion and/or deletions. Accuracy based on our ground truth set was between 80-90% for coding and 50-70% for protein variants, numbers that are not adequate for clinical reporting. Exact concordance for SNV syntax was over 99.5% between ClinVar and Variant Effect Predictor (VEP) and SnpEff, but less than 90% for non-SNV variants. For COSMIC, exact concordance for coding and protein SNVs were between 65 and 88%, and less than 15% for insertions. Across the tools and datasets, there was a wide range of equivalent expressions describing protein variants.

**Conclusion:** Our results reveal significant inconsistency in variant representation across tools and databases. These results highlight the urgent need for the adoption and adherence to uniform standards in variant annotation, with consistent reporting on the genomic reference, to enable accurate and efficient data-driven clinical care.

## INTRODUCTION

High-throughput sequencing has transformed the landscape of clinical genetic testing. This strategy, combined with the completion of massive public profiling datasets (ExAc [1], 1000 Genomes [2]), has dramatically changed our approach towards cancer treatment and the diagnosis of inherited disease. A major challenge in the analysis of this throughput and volume of data is integrating variant level information from the wealth of clinical and biological insight accumulated over decades of research, particularly those from recent, large sequencing studies. Describing a variant’s location is a fundamental part of a clinical assessment, yet the practice remains difficult, inconsistent and evolving.

Specifically, the clinical genomics community faces an enormous hurdle, which is integrating data generated prior to the availability of a robust human reference assembly with that generated using modern methods. Standards and guidelines for describing variants at the genomic, transcript (coding) and protein level, provided by the Human Genome Variation Society (HGVS) [3], were developed when testing was largely transcript rather than genome-based. As laboratories shifted to high-throughput sequencing, variant analysis transitioned to the genome level, confounding comparisons with reports generated from previous transcript-based assays.

Reconciling variant coordinates from the transcript to the genome, and vice versa, is not an unambiguous task. Requisite information about the genomic and transcript sequence accessions, their versions, and the alignments used to relate the two sequences, are not always reported in publications (Figure 1a–b). Alignment of cDNA to the genome remains challenging and can result in substantially different exon structures depending on the alignment approach (Figure 1a) [4,5]. In addition, variant reporting standards for VCF, a format designed to store genomic variation, are different from those for HGVS, a format that describes transcript and protein variants. In the context of nucleotide repeats, VCF shifts left with respect to the genome, while HGVS shifts right with respect to the gene or transcript (Figure 1c). Variants can therefore have very different locations depending on their accession, version and alignment.

**Figure 1.**
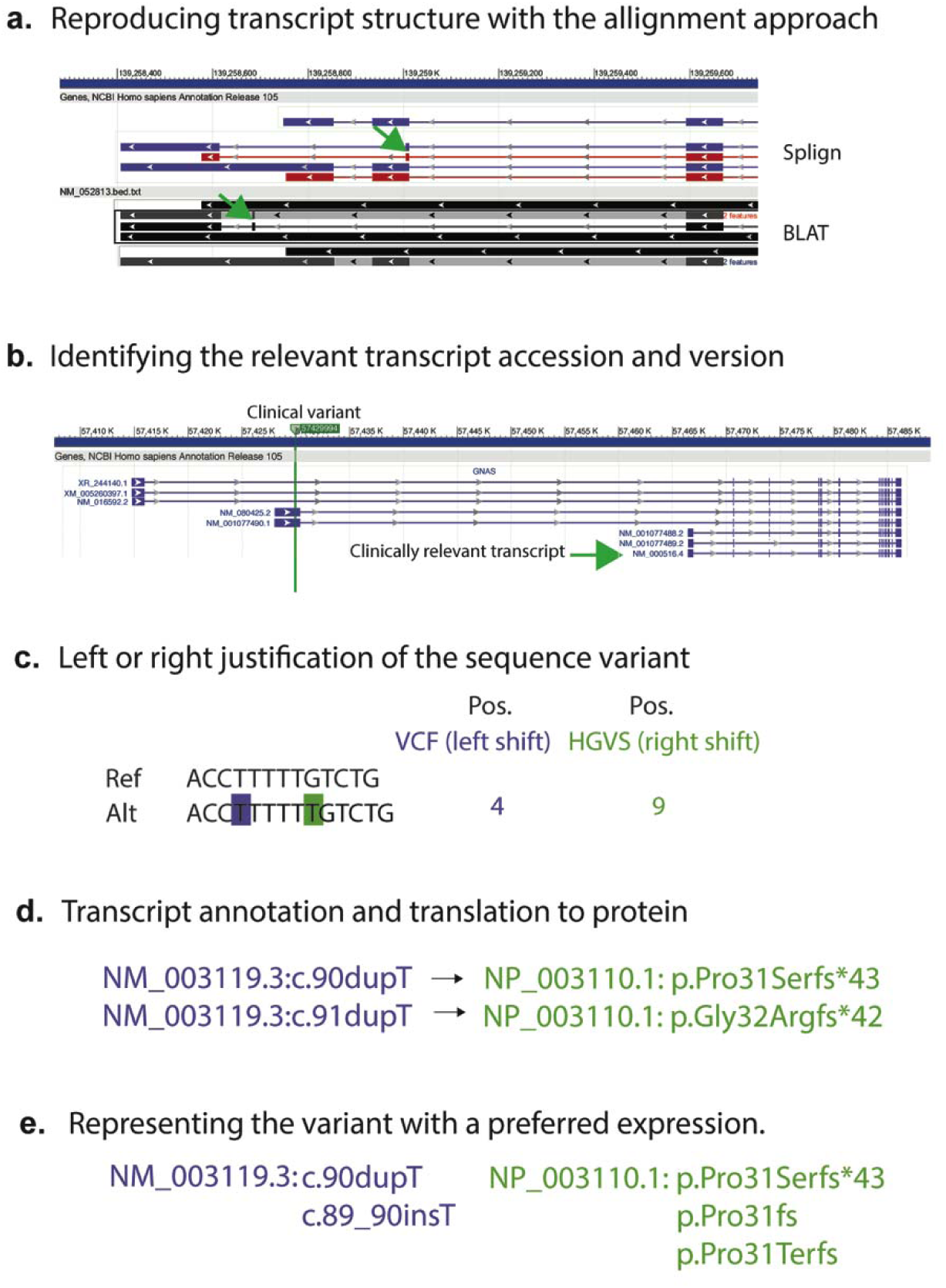
Factors affecting HGVS syntax generation. a) Transcript alignment approach can impact the transcript exon structure. Alignment of cDNA sequence by Splign and BLAT to the genome results in a 10kb difference in an exon positioning in the CARD9 gene (green arrow). b) Transcript accession can impact the variant association and HGVS syntax. Here, the identified GNAS variant is outside the clinically relevant transcript. Small changes in versions may also impact the coding sequence. c) In the context of nucleotide repeats, variant justification can affect the variant’s position. d) Transcript annotation directly impacts its translation to a protein expression. Incorrect transcript annotation can lead to incorrect protein syntax. e) Representing the variant in a particular expression. There are different ways of expressing the same coding or protein variant.

Even in relation to the same transcript, a variant can have multiple representations. HGVS expressions can have long and short forms, preferred and non-preferred syntax, and describe amino acids by their triple (e.g. Glu) or a single letter designation (e.g. E) (Figure 1d). In a survey by Deans et al. (2016) [6], 20 laboratories reported the HGVS syntax for a single variant in 14 different ways. An evaluation of over 140 molecular pathology laboratories in Europe and the UK revealed substantial errors in reported HGVS variant descriptions for the EGFR gene [7].

Currently there are many tools that automatically generate HGVS syntax, including SnpEff [8], Variant Effect Predictor (VEP) [9], Annovar [10], Variation Reporter (VR) [11], Mutalyzer [12] and packages developed by individual clinical laboratories such as Invitae [5] and Counsyl [13]. While the performance of different genomic variant callers have been well-studied [14,15], the accuracy and consistency of HGVS generation tools have not yet been described.

Previous comparison of Annovar and VEP revealed substantial differences in annotation based on choice of transcript [16]. This low concordance, combined with the increasing demand for automated syntax generation, prompted our re-evaluation of the performance of well-supported, open source tools. We considered only freely available tools as they would have the largest reach. Additionally, we wished to focus on annotation differences that can occur even when the same transcript is used. In this paper, we compare the concordance of variant nomenclature generated by VEP [9], SnpEff [8] and Variation Reporter, benchmarked by a curated ‘truth’ set and variant annotations described in large public datasets for germline (ClinVar) and cancer (COSMIC) variant descriptions. We find that while the tools SnpEff and Variant Effect Predictor produce comparable results, there remains significant discordance in variant annotation among tools, public resources, and literature.

## METHODS

### Datasets

We curated a test set of 127 variants to establish a ground-truth set with which we can evaluate the accuracy of the tools. Fifty-one variants were selected from public repositories: ClinVar, dbSNP, COSMIC, My Cancer Genome, Emory database and Leiden (Additional file 1: Table S1). We added 76 synthetic variants to ensure representation across variant types and genomic features. Genomic, coding and protein nomenclature for all variants were generated using a combination of the Mutalyzer [17] and Variation Viewer [18] webservice. Effect impact was determined based on the protein syntax and sequence ontology (SO) [19].

We used the ClinVar GRCh37 VCF and annotations from the tab separated file downloaded from the FTP site [20] (January 5^th^ 2016 release). We used the rsid and alternative allele to connect variants between the two files. We obtained genomic coordinates from the COSMIC GRCh37 VCF and connected them with the transcript and nomenclature in the CosmicCompleteExport.tsv file from the COSMIC website [21] (v75) with the COSMID.

### VCF normalization

We used vt-normalize [22] to left-justify all variants in each of the dataset VCFs used. A breakdown of insertions and deletions (indels) for each dataset and the number normalized is represented in Additional file 1: Table S2.

### Tools used

We ran SnpEff (v4.1L) [8,23], VEP (v82) [9] and VR [11] on our ground truth set, and subsequently only SnpEff and VEP on the ClinVar and COSMIC datasets. The Snpeff database was built using the NCBI GRCh37 GFF corresponding to the NCBI annotation ‘*Homo sapiens* 105’. The Snpeff database for Ensembl transcripts was built using the GRCh37 Ensembl transcript GFF [24]. We ran VEP with the corresponding RefSeq or Ensembl cache (v83). For all tools we used NCBI GRCh37p13 as the input reference genome.

### Assessment of syntax

To assess the performance of the variant annotation tools, we performed string match comparisons between the output and the reference syntax (Figure 2). Annotations were evaluated according to the HGVS guidelines [25, 26]. Variant annotations were labeled as ‘exact’ matches when the HGVS string and the query annotation matched as-is. If the string did not match perfectly, but could be transformed to the query string by applying HGVS recommendations, the tool’s annotation was labeled ‘equivalent’. For this study, both ‘exact’ and ‘equivalent’ annotations are regarded as correct. The code and module for performing the syntax assessment will be distributed on GitHub.

**Figure 2.**
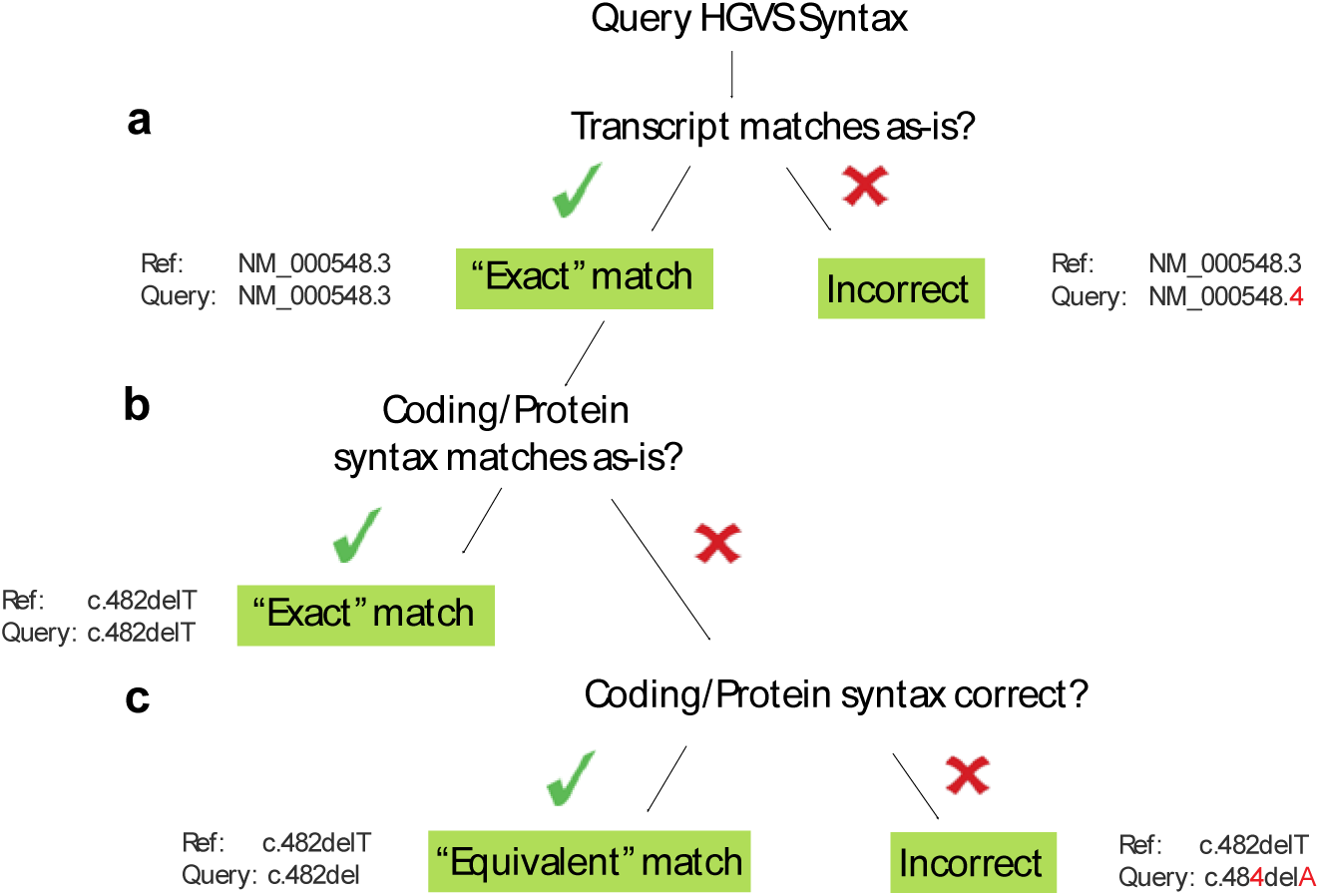
Methodology of HGVS syntax comparison. To compare two HGVS expressions in our dataset, we applied the following assessments. a) The query transcript must match the reference transcript. If the accession or version does not match, the variant is not assessed. b) If the syntax for both expressions correspond as-is, the match is ‘exact’. c) If the syntax for both expressions are equivalent, the match is ‘equivalent’, If the syntax is not an alternative expression of the other HGVS variant, the match is ‘incorrect’.

## RESULTS

### Comparisons to a ground truth test set

In order to assess the performance of different variant annotation tools against a ground truth, we used a contrived test set of 127 manually curated variants (Figure 3a) comprised of 52 previously reported variants in the literature or databases, and an additional 76 synthetic variants targeting a spectrum of variants (Additional file 1: Table S3). All annotations were reviewed manually using a combination of the Mutalyzer and Variation Viewer web services. This structured test set would allow us to deeply evaluate variants across different classes, effects, and genomic features.

**Figure 3.**
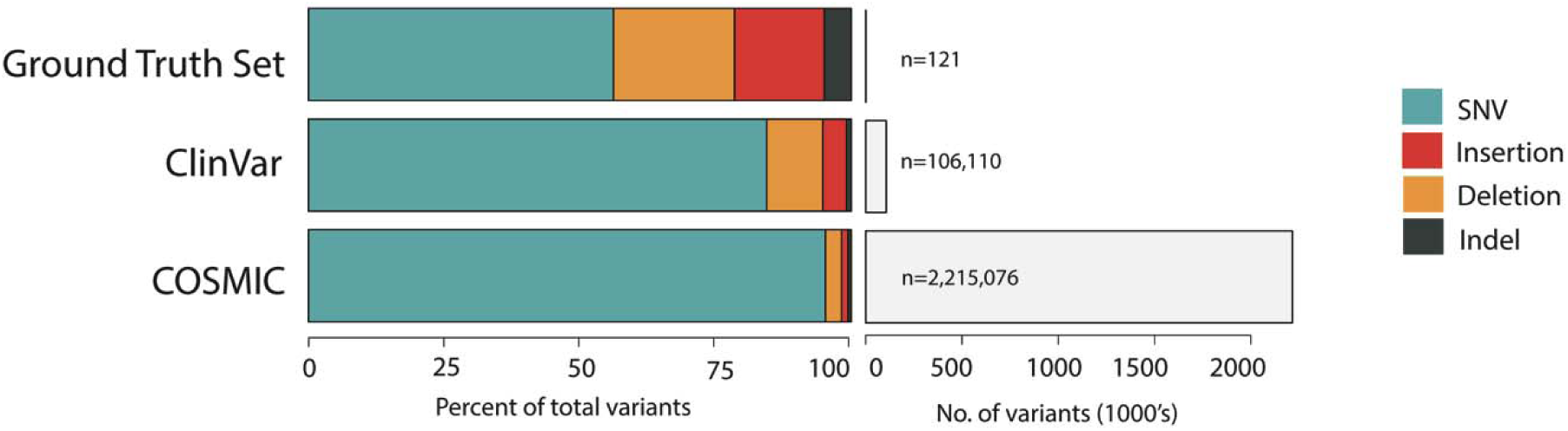
Datasets by composition. a) Number of variants evaluated in the Ground Truth, ClinVar and COSMIC dataset. Note that the number of variants assessed may be less than the number of variants in the input set. b) Distribution of variant types for each dataset. Duplications are included under insertions.

Using the analysis flowchart summarized in Figure 4, we compared the annotations generated by VR, VEP [9] and SnpEff [8] to the ground-truth test set (Additional File 1: Table S4). VEP and SnpEff accept VCF as input files; at the time of analysis, the VR API was limited in its functionality in processing large VCF files. Input HGVS expressions were also required for Mutalyzer, but we did not assess this tool because it was used to construct the ground truth set. We compared only annotations made on the same RefSeq transcript version. Although the input transcript alignments for SnpEff and VEP were identical, the tools produced a different number of transcripts and annotations. For example, we could not extract the relevant transcript for 4 variants in the SnpEff output and 5 from the VEP output, in addition to 5 variants absent from both tools. The importance of transcript collection was more pronounced for VR, which uses its own in-house alignments. As a result, 18% of the test variants could not be assessed by VR because NCBI carries only the most up-to-date transcripts. VR also frequently yielded multiple annotations for a single variant and transcript. In these cases, we chose the first variant in the output to evaluate in this test. In total, only 121 out of the 127 variants were annotated on the relevant transcript for any of the three tools.

**Figure 4.**
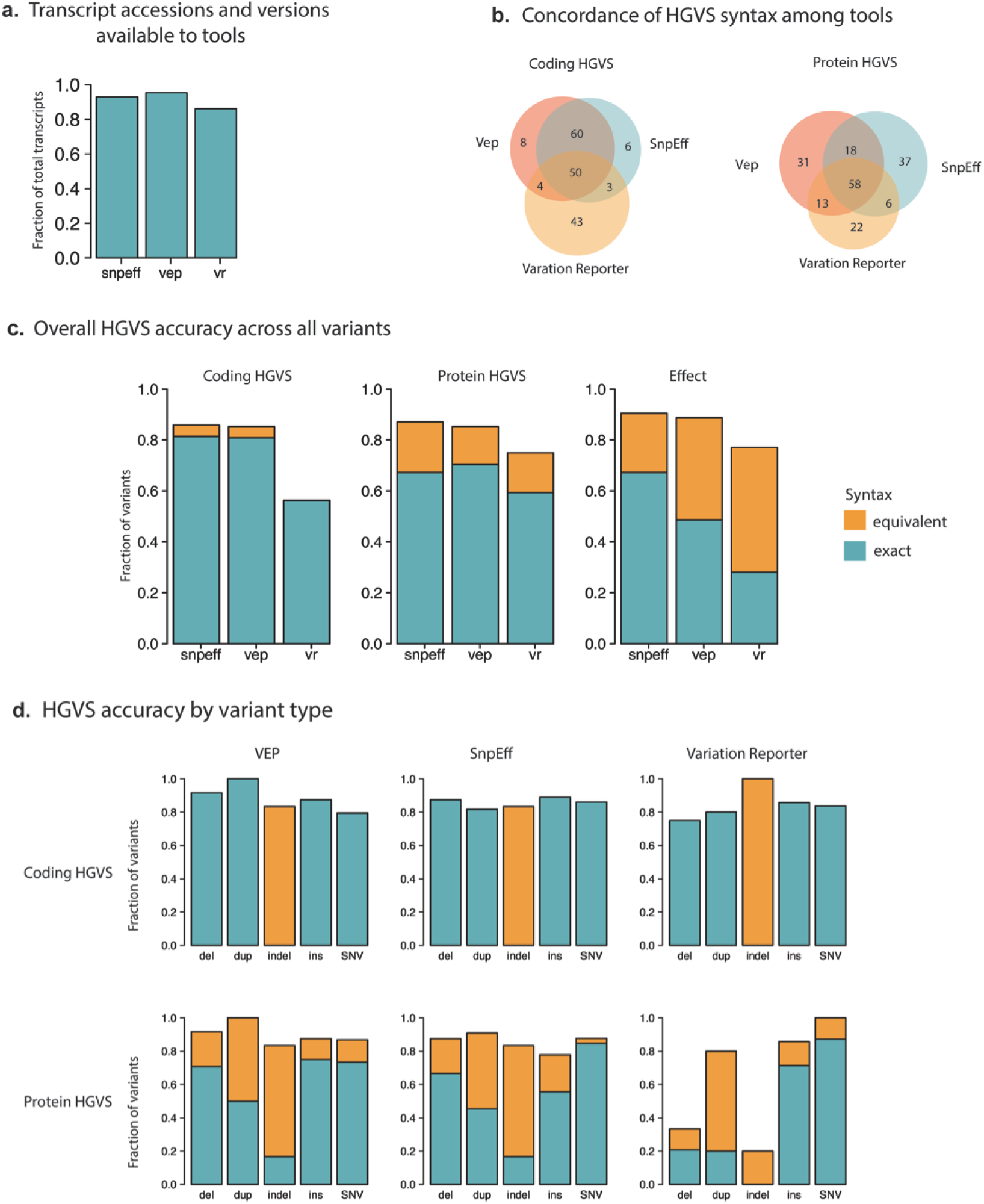
Summary of ground truth set HGVS syntax assessment. a) Fraction of unique transcript accessions and versions in the ground truth set that were available to the tools SnpEff (snpeff), VEP (vep), and Variation Reporter (vr). If a transcript was not accessible to the tool, the variant could not be annotated with respect to that transcript. b) Exact concordance of HGVS syntax at the coding (left) and protein (right) level among the tools. c) Accuracy of annotation across variants (n=121) described as exact (blue) and equivalent (orange). Fraction shown is with respect to number of annotations on the relevant transcript on the test set. d) Accuracy of annotation for each variant type across the tools. Variant types evaluated were: deletions (del), indels (delins), duplications (dup), insertions (ins) and single nucleotide variatnts (SNVs).

A major challenge in comparing nomenclature between tools was evaluating the equivalency of the many HGVS expressions for a given variant. Protein variant syntax was far more variable than coding variant syntax: between 14-20% of protein annotations were described with equivalent nomenclature across the three tools, compared to only 2.5% to 3.0% of coding syntax (Figure 4c–d). Each tool had distinct frameshift and synonymous annotations; frameshift has both long and short form alternatives, while synonymous variants can be described in several ways; e.g. ‘p.(=)’, ‘p.=’, ‘p.Thre258=’, ‘p.Thr258Thr’ (PTV012, Additional file 1: Table S5). Exact concordance in annotation between SnpEff and VEP was higher at the coding level (77.6% of variants) than at the protein level (68.8.1%) (Figure 4b, right panel). Less agreement was observed between Variation Reporter and either VEP or SnpEff: approximately 50% with either tool for coding and protein syntax.

For variants in our ground-truth set, SnpEff and VEP exhibited comparable accuracy and precision. At the coding level, SnpEff, VEP and VR annotated between 80% and 85% of substitutions correctly (out of 65 and 68), compared to 100% of substitutions for VR (out of 55). For deletions and insertions, VR performed poorly largely due to systematic errors in reporting. VR incorrectly described all but two deletions as indels. The remaining two annotations diverged from HGVS guidelines by omitting the ‘del’ designation altogether (e.g. c.2199-1301GA>A) (PTV062, PTV067, Additional file 1: Table S5). Duplications were also annotated as indels, but with technically equivalent (and redundant) nomenclature (c.1961dupG as c.1960delCinsCG). Such VR errors at the coding level led to inaccurate protein syntax for 18 variants.

We tested the ability of the tools to discriminate between the genomic reference and RefSeq transcript sequences, both of which are independently curated by the NCBI [27]. Since RefSeq transcripts typically receive a high level of manual review, conflicts between the RefSeq and genomic sequence often reveal an error in the latter. For this reason, we included nine test instances of RefSeq-Genomic differences in our ground truth set. Strikingly, none of the nine test examples of RefSeq-Genomic differences were identified by either VEP or SnpEff (Additional file 1: Table S5), and were erroneously reported as missense or deletion variants. While VR correctly identified 7 out of 9 RefSeq Genomic differences (the remaining two variants were not annotated), it mistakenly called differences for an additional 22 variants, indicating a poor precision for recognizing true differences. HGVS expressions should always reflect the base on the relevant genomic or transcript sequence to avoid asserting variants at positions where there is no change.

Both SnpEff and VEP correctly annotated the phased dinucleotide substitutions, which are variants present in consecutive bases, also known as multinucleotide variants (MNV) (Additional file 1: Table S6). Dinucleotide substitutions are highly prevalent in cancers associated with clear mutagen exposures such as melanoma, lung adenoma and lung squamous cell carcinoma [28]. Similarly, treatment by the chemotherapeutic agents cisplatin and meclorethamine have also been shown to cause dinucleotide substitutions at appreciable rates [28]. VR incorrectly annotated the phased dinucleotide substitutions as frameshift variants (PTV044, PTV068). We found that MNVs must be phased in the VCF, as the tools annotated adjacent but independent substitutions in the VCF separately instead of as a pair. For example, two BRAF variants (PTV045, PTV046) were incorrectly annotated as p.V600E and p.V600M, when the combined result would be p.V600K. These results indicate that for cancers with a high mutation load, prior phasing for dinucleotide pairs will be especially crucial to circumvent potential clinical oversights [29].

To complement the analysis of protein and coding annotations, we also assessed the variant effects predicted by the tools. Predicted effect is commonly used for evaluating pathogenicity during variant interpretation [30]. In instances where a variant could be associated with two functional consequences (for example, as intronic but also at a slice acceptor site), the annotation was considered to be correct if one association was described. Overall, the accuracy of effect prediction correlated highly with that of protein annotation (Additional file 1: Figure S1) even if they are calculated independently [8]). Compared to coding and protein syntax, efforts among tools to converge on a standardized set of variant effect annotations were far more evident (Additional file 2: Figure S1).

### Comparison with the ClinVar dataset

Having established baseline accuracy for automated syntax generation, we sought to assess the syntax concordance of these tools with those in public datasets. We started with ClinVar [31], a large public archive of variant and disease relationships that is widely used for evaluating Mendelian disease. Of the 106,110 small variants in the ClinVar VCF, the vast majority are SNVs (84%); the rest comprise a smaller number of deletions (10%), duplications (3.3%), insertions (1%) and indels (1%) (Figure 3b). We evaluated the performance of VEP and SnpEff on the ClinVar dataset (Additional file 3); because of the limited functionality and long running time of the VR tool, we did not include it in subsequent annotation assessments (Additional file 1: Table S7).

Approximately 10% of transcripts in the ClinVar dataset had different versions from those in our input transcript alignment file, which was used to build resources for both VEP and SnpEff (Figure 5a). Approximately 1.8% of ClinVar transcript accessions were not represented in the alignment input at all. Because of these discrepancies in transcript accession and versions, we could not assess the SnpEff or VEP annotations for 7 and 7.5% of ClinVar variants, again underscoring the importance of the input transcript set.

**Figure 5.**
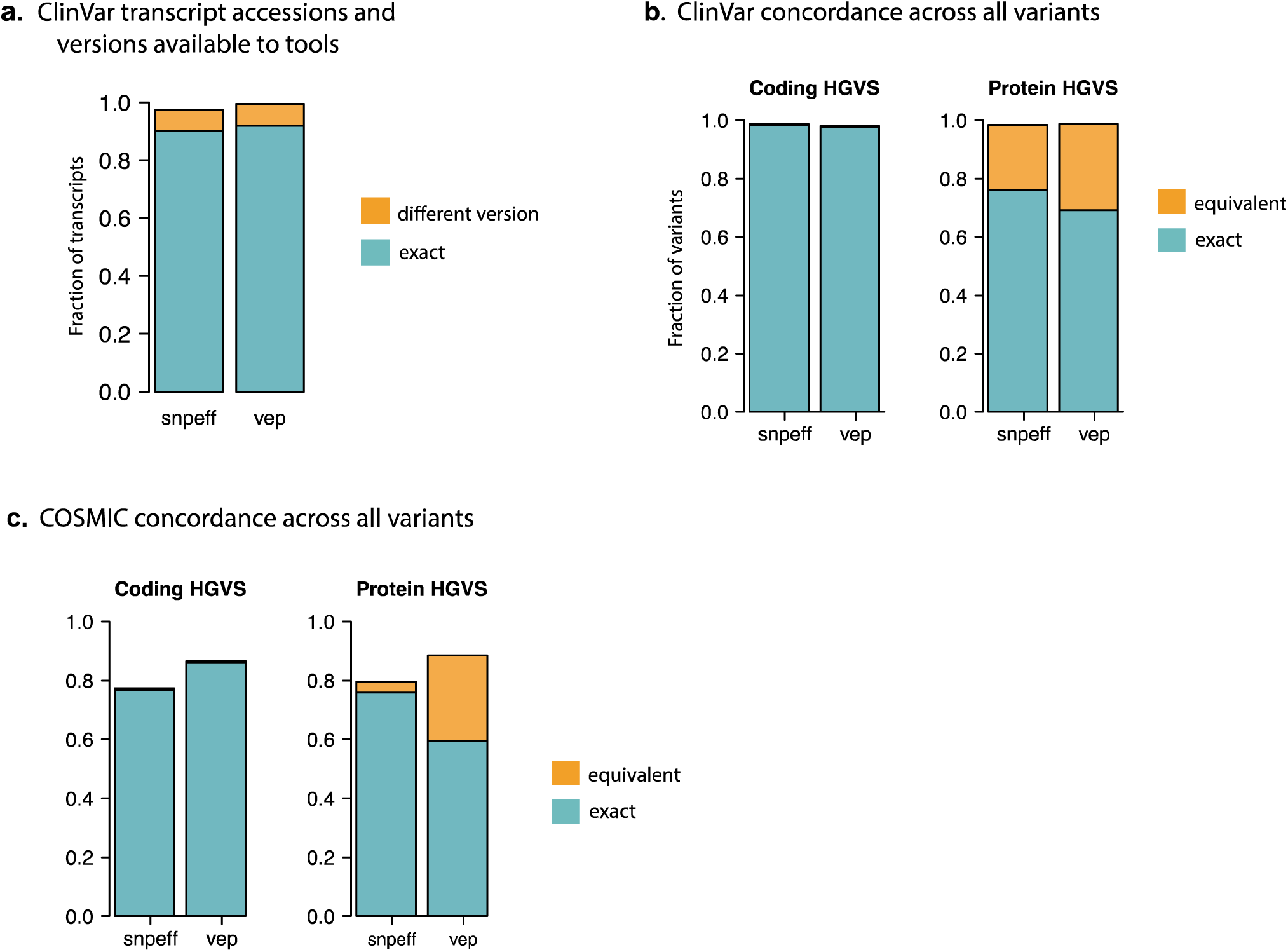
ClinVar and COSMIC HGVS syntax assessment. a) ClinVar transcript accessions and versions available to tools. Transcripts available to tools that matched the ClinVar reference transcript are marked in blue; transcripts that were different versions from the ClinVar transcript are marked in orange. Only unique transcripts were considered. b) Overall concordance in variant syntax across all variants between tools and ClinVar at the coding (upper panel) and protein (lower panel) level. Bars represent fraction of exact (blue) and equivalent (orange) matches. c) Overall concordance in variant syntax across all variants between tools and COSMIC at the coding (upper panel) and protein (lower panel) level. Bars represent fraction of exact (blue) and equivalent (orange) matches. All duplications were considered insertions in COSMIC.

Overall concordance for both SnpEff and VEP was remarkably high, which can be attributed to the large proportion of SNVs (Figure 5b). At the coding level, both SnpEff and VEP yielded nearly perfect concordance for SNVs, matching the exact ClinVar nomenclature for over 99.9% of the SNVs (Figure 5a, Additional file 2: Figure S2b). In rare instances of error, SnpEff and VEP were typically incorrect by one base (Additional file 1: Table S8). Exact concordance was lowest for variants annotated as insertions in ClinVar (approximately 75-80% for both tools), largely due to their correct assertion as duplications by VEP and SnpEff. In contrast, concordance was slightly higher for deletions and indels (between 86 and 88%). There were 25 instances in which neither tool could have predicted correct coding HGVS syntax without prior reports of the splicing product. A single nucleotide change at a splice site in the AGA gene NC_000004.11:g.178354367C>A (NM_000027.3:c.940+1G>T) results in the skipping of exon 8 and a final syntax of c.807_940del134 (Additional file 1: Table S9). In five of these cases, this type of error also resulted in the incorrect protein syntax.

As with the ground truth test set, we observed greater variation in protein syntax (Table 1). This was mostly evident for deletions, duplications, and insertions, where between 16 to 78% of annotations were reported correctly but with alternative nomenclature (Additional file 2: Figure S2b). Overall concordance was again high for SNVs (99%), with 75% and 83% exact nomenclature for SnpEff and VEP. However, neither VEP nor SnpEff performed as well on deletions, duplications and insertions (between 76.3% and 94.4% overall concordance). For non-SNV variant types, our results show that between 60-70% of annotations output by these tools do not match the ClinVar HGVS, and between 5-20% of these annotations are completely discordant. Dinucleotide substitutions, which ClinVar reports as indels, were annotated as independent substitutions for both SnpEff and VEP (Additional file 1: Table S6). In a few cases at the boundaries between coding and non-coding regions, VEP and ClinVar yielded no output while SnpEff reported the ambiguity as ‘p.Thr662_Glu663delins???’. Even for substitutions, there were instances where all three tools yielded distinct nomenclature for the same variant. For NM_001126128.1:c.163delA, ClinVar, SnpEff and VEP output p.Ile55Terfs, p.Ile55fs, and p.Ile55Ter respectively. The correct HGVS syntax for this variant is p.lle55Ter.

**Table 1.**
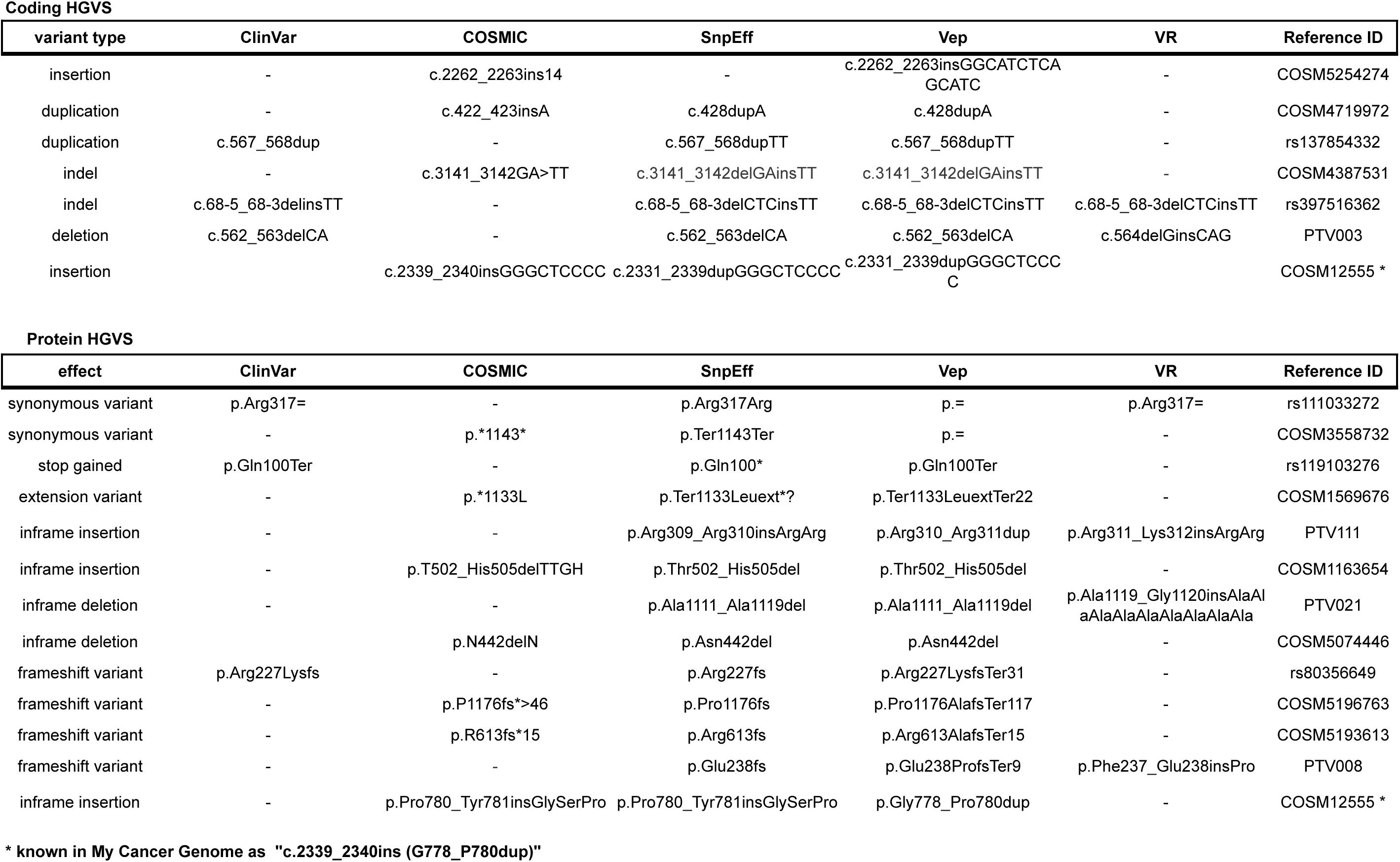
Exemplar variants demonstrating nomenclature discrepancies

Interestingly, agreement between SnpEff and VEP sometimes revealed errors or inconsistencies in the ClinVar output. For example, for rs34618570, the TTN variant NM_133378.4:c.10361-2293A>T is purported to be a missense variant (p.Ile3877Phe), when the variant is intronic for that transcript. For rs398123611 (Additional file 1: Table S8), ClinVar recognizes NM_133378.4:c.1138_1140dupGGC as a duplication in the coding syntax but annotates the protein as an insertion (p.Gly380_Ala381insGly). At least 626 variants in the ClinVar dataset were incorrectly annotated as both nonsense and frameshift (p.Glu307Terfs), when the output should simply be nonsense. Together, these results demonstrate that while there is near perfect consensus between the HGVS tools and ClinVar annotations for SNVs, the uniformity and correctness for other variant types, which are often of the most clinically relevant (e.g. frameshifts), still needs improvement.

### Comparison with the COSMIC dataset

Clinical cancer care is dependent on identifying relationships between tumor variants and relevant information about their prognostic and therapeutic significance. We investigated the consistency between annotation output by SnpEff and VEP with COSMIC, currently the largest public resource of somatic mutations in human cancer [32] that is also widely used by clinical laboratories. Again, we did not include VR in our assessment because of its limited functionality and long running time. Because COSMIC annotates variants in relation to Ensembl instead of NCBI RefSeq transcript accessions, we built a second, separate database to run VEP and SnpEff according to Ensembl transcript alignments.

We queried a total of 3,075,504 coding COSMIC variants. Following normalization and de-duplication of the COSMIC VCF, there remained a set of 2,215,076 variants (Figure 3). Approximately 142,134 variants were insertions, deletions or indels, 19% of which required left justification (Additional file 1: Table S2). We compared syntax representations (Figure 5c, Additional file 4). Both SnpEff and VEP generated annotations for approximately 90% of the COSMIC dataset. Because the cancer field employs the convention of abbreviating amino acids to a single letter while the annotation tools, and HGVS, all use the three-letter convention, we converted the COSMIC annotations to three-letter amino acids to facilitate annotation comparison.

At the coding level, VEP recapitulated the exact syntax as COSMIC for 85.9% of the total variants, compared to 76.8% of variants by SnpEff, with less than 1% of equivalent syntax for both tools (Figure 5b). However, the majority of the COSMIC dataset are SNVs (95%); for variant types other than SNVs, neither VEP nor SnpEff achieved comparable concordance (Additional File 2: Figure S2b). Notable differences in annotations include COSMIC’s reporting of all duplications as insertions, resulting in nearly complete discordance for variants of this type. We did not assert the equivalency of multi-base insertions with duplications due to the involvement of verifying duplicated bases in the reference transcript. As a result, none of the indel annotations were exact string matches. Additionally, in complete departure from current HGVS standard, COSMIC reports indels as block substitutions (c.569_570TC>AT vs c.569_570delTCinsAT, Table 1), which we assessed as ‘equivalent’. This format could be attributed to the historical representation of dinucleotide variants [33], which remains popular despite the adoption of the HGVS standard by most clinical resources (e.g. My Cancer Genome). By failing to consistently right justify insertion and deletion positions, the concordance between tools and COSMIC nomenclature for deletions was less than 50% (Additional file 1: Table S10).

For protein variants, SnpEff reproduced the exact protein syntax for 75.8% of COSMIC variants compared to 58.4% by VEP (Figure 5b). A large fraction of VEP discordance could be attributed to VEP’s annotation of all frameshifting indels as nonsense variants (Additional file 1: Table S10, COSM1476431). Further, over 90% of VEP alternative protein expressions were due to discrepant reporting of synonymous variants as p.= compared to p.Gly35Gly by both COSMIC and SnpEff (Table 1). Similar to coding deletions, nuances in nomenclature revealed distinct expressions of frameshifts for COSMIC, VEP and SnpEff.

For the majority of discordant annotations, the agreement between SnpEff and VEP syntax suggest that the COSMIC syntax is incorrect. To verify the HGVS nomenclature of these variants, we mapped the Ensembl transcript to its approximate corresponding RefSeq accession through its consensus coding sequence (CCDS), since a number of tools, including Mutalyzer, do not support Ensembl identifiers. A mutation in *TP53* at position chr17:7578525 (COSM1683507) is annotated in COSMIC as c.404_405insC. Because of a sequence of 4 C’s at this position, the standardized left shifted VCF position should be at chr17:7578523 and right-shifted HGVS syntax as c.405_406insC, or c.405dupC. In another example, a HER2 insertion variant is described in My Cancer Genome as c.2339_2340ins (with no insertion bases or transcript as reference) and G778_P780dup. The correct coding syntax by both SnpEff and VEP is c.2331_2339dup while the correct protein syntax (output only by VEP) is p.Gly778_Pro780dup. COSMIC annotated neither the coding or protein syntax correctly (Table 1). Based on the agreement of VEP and SnpEff alone, our results suggest that at least 2.7% of COSMIC variant annotations are incorrect (Table 1). This is not surprising given its recent transition from a research repository to a major clinical resource, although efforts to comply with genomic and HGVS standards are apparently underway.

### Clinical impact of discordant variant annotation

Ultimately, we are concerned about the concordance of positional and syntax expressions because of its impact on clinical interpretation. To illustrate this point, we describe a frameshift variant in the *PROK2* gene, which was differentially classified as an exercise by two curators in our laboratory - one classifying as likely pathogenic and the other as pathogenic for Kallman syndrome. The difference in classification stemmed from the use of different syntax in constructing the string-based search. The variant was described as ‘NM_001126128.1:c.297dupT (p.Gly100Trpfs*22)’. Because of alternative transcripts and HGVS representations, this variant could be searched by multiple expressions (Additional file 1: Figure S3a). In one route, searching ‘PROK2 c.297_298insT’ or ‘PROK2 c.234_235insT’ immediately retrieved the relevant literature to classify this variant. However, searching ‘PROK2 *297_298ins*’, ‘PROK2 *234_235ins*’, or the correct HGVS syntax ‘c.297dup’ or ‘c.234dup’ did not return any relevant results (Additional file 1: Figures S2b). Searching for ‘PROK2 G100fsX121’, ‘PROK2 c.297_298insT’ or ‘PROK2 c.234_235insT’ identifies a paper by Abreu et al. [34], which leads to a thread of reports that supports a final variant classification of ‘pathogenic’ (Additional file 1: Figure S3b-c). Because of these multiple variant representations, identifying relevant information can entail navigating a complex matrix of HGVS expressions and web results.

As another example of the importance of accurate HGVS nomenclature for clinical care, a variant in a patient’s melanoma sample was annotated in our pipeline as ‘NM_004333.4:c.1799T>A (p.V600E)’. During visual review we found that the variant was part of a dinucleotide pair, with a combined syntax of c.1799_1800delTGinsAT and protein syntax of p.V600D. Although p.V600D is sensitive to BRAF inhibitors, this variant is not as well-studied and characterized with respect to drug response and efficacy compared p.V600E. Further, while V600E confers sensitivity to MEK inhibition, the sensitivity of p.V600D to MEK remains unclear.

## DISCUSSION

We have described some of the remaining challenges of moving clinical sequencing into a high-throughput environment. Consistent with findings by McCarthy et al. [16], we find that the transcript collection has a significant impact on the yield of relevant variant annotations. Our examination of automated syntax from HGVS tools and the ClinVar or ground truth datasets reveal that approximately 10% of variants could not be assessed due to discordant transcript accessions or versions. The fact that ClinVar and COSMIC, the largest public repositories of germline and somatic data respectively, do not share the same collection of transcript accessions reflects the degree of harmonization and the need for a universal store of transcript to genome alignments.

Importantly, although variant calling is performed almost exclusively on genomic data, variants are still being primarily referenced with respect to the transcript. Recent publications continue to describe variants according to their protein and/or coding syntax [35–37], sometimes even without the transcript identifier [38,39]. In a survey by the American Society of Molecular Pathologists, 50% of clinical cancer labs report variants exclusively by coding and protein HGVS nomenclature but without accompanying genomic coordinates. The same survey also found that 70% of clinical cancer labs use as a resource MyCancerGenome.org, which references variants by their popular single-letter amino acid or coding-level convention, again, without transcript or genomic coordinates. As our analyses show, transforming genomic positions to transcript loci is challenging and prone to error; ambiguity in representation is best avoided by always referencing variants by their genomic position and assembly version. For this reason, HGVS recommends reporting clinical variants by their Locus Reference Genomic sequence (LRG), a system designed for clinically relevant variants that is based on un-versioned and stabled accession sequences [26,40].

Despite the precision achieved with generating syntax for SNVs, the positions of insertions and/or deletions remain stubbornly difficult to annotate, regardless of the VCF or HGVS genomic standard. The presence of duplicates in nearly one-fifth of the COSMIC VCF highlights the importance of using tools for normalization to reconcile the multiple possible positions to represent a single variant. At the level of HGVS, we found that none of the non-SNV variant types were annotated with near 100% accuracy or compliance with HGVS conventions for any of the tools or databases that we queried. Given the rigorous reporting requirements of a clinical genetics lab, this is concerning, and suggests that it remains critical to manually review the syntax when reporting non-SNVs.

Our analyses further provide a glimpse into the diverse matrix of possible HGVS representations for a given variant - a disturbing concept for attempts to mine and exploit existing resources through string-based search. Internal efforts can be made to standardize HGVS syntax within knowledge-bases and clinical enterprises; variants can be transformed into a standard, minimal expression to enable a uniform query across curated databases [41]. However, while this is useful for a limited set of data, it is impractical for mining beyond internally curated information. The alternative is exhaustive but impractical, requiring the search for every permutation of an HGVS expression for a particular variant. A thoughtful discussion should be made about asserting HGVS guidelines as rules to enforce a strict convergence across laboratories, resources, and literature.

By design, the HGVS annotation system was not intended for mining large bodies of genomic information, while approximations of syntax are not acceptable because of their impact on clinical care. A means of clinical intervention in oncology is to directly connect clinically actionable variants in patient tumor samples with relevant therapeutic strategies, such as approved drugs or eligibility for clinical trials. In the ACMG guidelines for the classification of germline variants, at least five categories of evidence require interrogating variants from previous reports in reliable databases or the published literature [30]. Already, studies have shown that there remains substantial heterogeneity in the interpretation of genomic variants by clinical laboratories [6,7,42]. Imprecise nomenclature can lead to variant misclassification and consequent misdiagnosis [29]. The applications of genomics in clinical care will require concerted efforts to converge on standardized reporting mechanisms to enable data sharing and integration across diverse datasets and resources. Reporting on the same genomic reference, according to uniform variant syntax, will be one crucial step towards the achieving this aim and the ultimate goal of precision medicine.

## DECLARATIONS

### List of abbreviations

HGVS, Human Genome Variation Society; VEP, Variation Effect Predictor; VR, Variation Reporter; VCF, Variant Call Format

### Competing Interests

JY, SG, AM, JH, SC, JW, RC, DMC were full time employees of Personalis at the time of this study.

### Authors’ contributions

JY designed the study, performed the bioinformatics and data analysis, interpreted results and wrote the manuscript. DMC conceived the idea and guided the study design, results interpretation and manuscript preparation. SG contributed to the data analysis, study design and results interpretation. AM and SC contributed to the bioinformatics analysis. Both AM and JH provided guidance for the bioinformatics analysis. All authors read and approved the final manuscript.

## Acknowledgements

We thank Dariya Glazer and Joshua Anderson for their contributions in classifying the PROK2 variant; Christian Haudenschild for his guidance on the dataset submission and Massimo Morra for his keen eye in HGVS nomenclature.

### ADDITIONAL FILES

**Additional file 1**

**Table S1**. Ground Truth Set Variants

**Table S2**. Variants Normalized by Dataset

**Table S3**. Ground Truth Set Contents by Features

**Table S4**. Ground Truth Set Comparison Results. Exact matches between the reference annotation in COSMIC and annotations provided by Snpeff and VEP are noted as “yes”, equivalent matches as “yes_m” (“yes modified”) and not equivalent annotations as “no”.

**Table S5**. Example Nomenclature Discrepancies Ground Truth Set Variants

**Table S6**. Annotation of dinucleotide substitutions

**Table S7**. Tools Run Time

**Table S8**. Example Nomenclature Discrepancies from the ClinVar Dataset

**Table S9**. Examples of genomic SNVs resulting in deletions at the transcript level

**Table S10**. Example Nomenclature Discrepancies from the COSMIC Dataset

**Additional file 2**.

**Figure S1. Comparison of effect annotation between tools and the HGVS test set**.

a) Concordance in effect nomenclature between the HGVS test set and SnpEff and VEP by variant type.

**Figure S2**. Concordance in variant syntax by variant type between tools and a) ClinVar or b) COSMIC datasets at the coding (upper panel) and protein (lower panel) level. Bars represent fraction of exact (blue) and equivalent (orange) matches. All duplications were marked as insertions in COSMIC.

**Figure S3. Impact of HGVS nomenclature on clinical interpretation**

a) Transcripts and nomenclature associated with variant (chr3:g.71821968dupA).

b) PubMed and Google results from search strings.

c) From a single search string to evidence and classification.

**Additional file 3**. ClinVar Comparison Results. Exact matches between the reference annotation in COSMIC and annotations provided by Snpeff and VEP are noted as “yes”, equivalent matches as “yes_m” (“yes modified”) and not equivalent annotations as “no”.

**Additional file 4**. COSMIC Comparison Results. Exact matches between the reference annotation in COSMIC and annotations provided by Snpeff and VEP are noted as “yes”, equivalent matches as “yes_m” (“yes modified”) and not equivalent annotations as “no”.

